# Profiling the organ membrane proteome dysregulation in the context of liver disease

**DOI:** 10.1101/2024.04.05.588300

**Authors:** Frank Antony, Zora Brough, Mona Orangi, Hiroyuki Aoki, Mohan Babu, Franck Duong van Hoa

## Abstract

Alcohol consumption and high-fat diets often coincide in Western society, exerting negative synergistic effects on the liver. While many studies have demonstrated the impact of ALD and NAFLD on organ protein expression, none have offered a comprehensive view of the dysregulation at the level of the membrane proteome. In this study, we utilize peptidisc and solvent precipitation (SP4) methods to isolate and compare the membrane protein content of the liver with its unique biological functions. Using mice treated with a high-fat diet and ethanol in drinking water, we identified 1,563 liver proteins, with 46% predicted to have a transmembrane segment. Among these, 106 integral membrane proteins are dysregulated compared to the untreated sample. Gene ontology analysis reveals several dysregulated membrane processes associated with lipid metabolism, cell adhesion, xenobiotic processing, and mitochondrial membrane formation. Pathways related to cholesterol and bile acid transport are also mutually affected, suggesting an adaptive mechanism to counter the steatosis of the liver model. Our peptidisc-based membrane proteome profiling thus emerges as an effective way to gain insights into the role of the transmembrane proteome in disease development, warranting further in-depth analysis of the individual effect of the identified dysregulated membrane proteins.

## 1. Introduction

Chronic liver diseases account for 2 million deaths annually, ranking as the eleventh leading cause of death worldwide [1]. Among the primary contributing factors are obesity and long-term alcohol abuse, both causing hepatic steatosis and inflammation. If untreated, the associated non-alcoholic fatty liver disease (NAFLD) and alcohol-induced liver disease (ALD) can progress to fibrosis, and eventually cirrhosis or hepatocellular carcinoma [2]. Since alcohol is primarily metabolized in the liver, where lipid processing also occurs, it is no surprise that its consumption combined with a high-fat (HF) diet synergistically accelerates liver injury and poorer outcomes [3-6].

The primary defect in chronic liver pathogenesis is dysregulated lipid metabolism and excessive accumulation of fatty acids and cholesterol within hepatocytes, known as hepatic lipotoxicity [7]. At the tissue level, these injuries promote inflammation, oxidative stress, apoptosis and extracellular matrix deposition among other deleterious effects [4, 8-10]. In the search for the molecular causes, several studies have observed the dysfunction of various membrane proteins (MPs). For instance, ethanol exposure and obesity were found to enhance the expression of fatty acid translocase CD36 [11,12] leading to heightened lipid accumulation within cells [13,14]. The activities of the endoplasmic reticulum Cyp P450 enzymes, monooxygenases crucial for cholesterol metabolism and detoxification of xenobiotics like alcohol [15,16], were also dysfunctional in ALD and NAFLD [17,18]. Similarly, the production cycles of bilirubin and reabsorption of bile acids, both profoundly affected by liver diseases, all depend on membrane importers and exporters [19-21]. Additionally, mitochondrial structural defects, especially in cristae formation, have been associated with liver dysfunction [22,23] whereas alcohol consumption affected inner membrane potential, activating caspase-9 and -3 pathways and cell apoptosis [24]. Another hallmark of liver diseases is a dramatic remodeling of the extracellular matrix, which involves cell-surface adhesion proteins such as CD44 that contribute to fibrosis progression and inflammatory responses [25-28]. From this, it is evident that ALD and NAFLD are linked to major alterations of the liver membrane proteome, and likely lipidome as well [29].

Several proteomic studies have been conducted on liver mouse models [30] but none has provided a comprehensive view of the regulation of the global membrane proteome. Studying MPs has proven challenging for multiple reasons. Firstly, MPs are underrepresented in the cellular pool and overshadowed by soluble proteins during mass spectrometry analysis [31]. This poses a particular challenge when studying liver tissue, which exhibits a dramatic proteome expression range, with the most abundant protein reported to be one million times more abundant than the least abundant ones [32]. Secondly, integral membrane proteins (IMPs) contain long hydrophobic segments that render them prone to aggregation, precipitating out of solution and becoming lost during sample preparation [33]. To address the challenge, here we evaluate the performance of two sample preparation techniques, namely the solvent precipitation SP4 and the peptidisc methods, in isolating the specificity of the liver membrane proteome. The method that best represents the liver function is then used to understand the dysregulation of IMPs in the context of liver disease. We report and document a specific set of membrane proteins whose dysregulation is related to liver malfunction.

### Significance of the Study

Liver diseases represent a concerning trend in chronic conditions, with rising mortality rates in recent years. ALD and NAFLD stand out as prevalent conditions in the Western world. Notably, the co-occurrence of alcohol consumption and high-fat diets has become increasingly common, exacerbating the risk of liver disease. To investigate the effect of the combined treatment, we chose to utilize a mouse model that mimics this scenario, exposing the mice to both an HF diet and ethanol consumption. The liver poses unique challenges for researchers due to its uneven distribution of protein abundances, especially integral membrane proteins (IMPs) that often linger in lower quantities relative to soluble proteins. Amidst this complexity, IMPs remain underrepresented in conventional proteomic methodologies despite their critical importance to the hepatocyte function. Hence, we directed our focus towards studying IMPs relative expression to gain deeper insights into liver pathophysiology.

## 2. MATERIALS AND METHODS

### 2.1 Materials

Detergents n-dodecyl-β-D-maltoside (DDM) and sodium deoxycholate (SDC) were purchased from Anatrace. His_6_-tagged peptidiscs (purity >90%) were obtained from Peptidisc Biotech. Ni^2+^-NTA chelating Sepharose resin was obtained from Qiagen. Silica beads and the complete protease inhibitor cocktail were purchased from Sigma. MS-grade trypsin and DNAase were purchased from Thermo Fisher Scientific. Octadecyl C18 Empore disks were purchased from 3M and Polygoprep 300-20 C18 powder was purchased from Macherey-Nagel. Centrifugal filters were from Amicon. All other general reagents such as NaCl, Tris-base, DTT, PMSF, iodoacetamide, urea, SDS, organic solvents and acids were obtained from Bioshop or Fisher Scientific Canada.

### 2.2 Mouse treatment and harvest of organs

The C57BL/6 mice were kept in specific pathogen-free conditions and received humane care in compliance with the Canadian Council of Animal Care guidelines, and the animal protocol was approved by the Animal Care Committee of the University of British Columbia. The mouse organs were obtained from six female mice. The authors acknowledge that they did not consider the impact of mouse sex at the time of the study design. At the age of 12 weeks, mice were randomly allocated into 2 experimental groups. One group (n=3) was fed with a diet made of 17% fat (50% lard and 50% cacao-butter, supplemented with 1.25% cholesterol and 0.5% cholate) plus alcohol in drinking water (1% for the first 2 days, 2% from day 3 to day 7, 4% in the second week, and eventually 5% final for another 7 weeks). The control group (n=3) received standard chow. After 9 weeks of feeding, the animals were sacrificed. The collected liver and lung organs were placed in vials with ice-cold sterile phosphate-buffered saline (PBS) until processing.

### 2.3 Tissue processing and preparation of the membrane fraction

The collected organs were rinsed five times in ice-cold PBS to remove blood, then minced and homogenized using a tight-fit metal douncer in a hypotonic lysis buffer (10 mM Tris-HCl pH 7.4; 30 mM NaCl, and 1 mM EDTA, 1x cocktail protease inhibitor and 1 mM PMSF). The homogenate was further incubated for 10 min on ice in the presence of 10 mM MgCl_2_ and 50 µg/ml DNAse. The enlarged cell suspension was lysed using a French press with 3 passages at 500 PSI. The cell lysate was centrifuged at 1,200 x g for 10 min at 4^◦^C to remove unbroken cells and nucleus fraction. The supernatant was collected and centrifuged at 5,000 x g for 10 min at 4^◦^C to remove the mitochondrial fraction and the supernatant was ultracentrifuged at 110,000 x g for 45 min at 4^◦^C in a Beckman TLA110 rotor. The pellet was resuspended in 200 μL of TSG buffer (50 mM Tris, pH 7.9, 100 mM NaCl, 10% glycerol) to obtain a membrane fraction named “crude membrane” and stored at -80°C until use.

### 2.4 Sample preparation for SP4-based analysis

The sample preparation was based on the protocol described by Johnston et al, 2021. Briefly, silica beads (9–13 μm diameter) were resuspended in water, washed once with 100% acetonitrile (ACN), rinsed twice with water and resuspended at 50 mg/ml final. After each wash, the beads were isolated by brief centrifugation at 16,000 x g for 1 min. Crude membranes (∼1 mg) were resuspended in ice-cold TS buffer (50 mM Tris-HCl, pH 8.0, 100 mM NaCl) supplemented with 1% (w/v) SDC for 30 min at 4^◦^C with gentle shaking. The detergent extract was clarified by ultracentrifugation (110,000 x g for 15 min at 4^◦^C) and aliquots (100 µg each) were gently vortex-mixed with glass beads (1 mg). Acetonitrile was then added to a final concentration of 80% and samples were centrifuged at 16,000 x g for 5 min. The beads were rinsed three times with 500 µl of 80% ethanol without disturbing the pellet. After a final wash, the beads were resuspended in 100 µl of 6 M urea and placed in a sonicator bath for 5 min. Proteins were reduced with 10 mM DTT for 1 hour, alkylated in the dark with 20 mM iodoacetamide and further reduced with 10 mM DTT for 30 min. The urea was diluted to 1 M with the TS buffer and trypsin was added at an enzyme/protein ratio of 1:100 for 24 h at 25°C on a shaker. The beads were recovered by centrifugation at 16,000 x g for 1 min and the supernatant (600 µL) was placed in a new tube. The samples were acidified with 10% formic acid to pH 3 (25 µL) and loaded onto hand-packed C18 Stage-tips. Samples were eluted with 80% ACN containing 0.1% formic acid. The eluted peptides were dried by vacuum centrifugation.

### 2.5 Sample preparation for peptidisc-based analysis

Crude membranes (∼1 mg) were resuspended in ice-cold TS buffer (50 mM Tris-HCl, pH 8.0, 100 mM NaCl) supplemented with 1% (w/v) DDM for 30 min at 4^◦^C with gentle shaking. The detergent extract was clarified by ultracentrifugation (110,000 x g for 15 min at 4^◦^C) and 1 mg aliquots (500 µl) were incubated with a 3-fold excess of His_6_-tagged peptidiscs for 15 min at 4^◦^C. The sample was rapidly diluted to 5 mL in the TS buffer placed in a 100 kDa cutoff centrifugal filter. The sample was concentrated at 3,000 x g, 10 minutes to approximately 200 µl and diluted again to 5 ml, followed by another concentration step to approximately 200 µl. The reconstituted start peptidisc library (4 mg total in 1 mL) was incubated with 60 μL of Ni^2+^-NTA chelating Sepharose for 1 hour at 4^◦^C with shaking. Following five thorough washes with TS buffer 1 mL each, the peptidisc library was eluted in 150 μL of TS buffer containing 600 mM imidazole, recovering 1 mg at a concentration of ∼6.5 µg/µL. All fractions from the reconstitution through the purification steps were subjected to analysis by 15% SDS PAGE, followed by Coomassie blue staining of the gel. An aliquot of the purified membrane protein library containing (∼100 μg; ∼16 µL) was treated with 6 M urea at room temperature for 30 min, followed by reduction with 10 mM DTT for 1 hour. Alkylation was performed with 20 mM iodoacetamide (IAA) in the dark at room temperature for 30 min, followed by a re-addition of 10 mM DTT for 30 min. The urea concentration was diluted to 1 M with the TS buffer and trypsin was added at an enzyme/protein ratio of 1:100 for 24 h at 25°C on a shaker. Subsequently, the digested peptides were acidified to pH 3 with the addition of ∼20µl 10% formic acid and desalted using hand-packed Stage-Tips C18. The eluted peptides were dried by vacuum centrifugation.

### 2.6 LC and MS/MS analyses

NanoLC connected to an Orbitrap Exploris mass spectrometer (Thermo Fisher Scientific) was used for the analysis of all samples. The peptide separation was carried out using a Proxeon EASY nLC 1200 System (Thermo Fisher Scientific) fitted with a custom-made C18 column (15 cm x 150 μm ID) packed with HxSil C18 3 μm Resin 100 Å (Hamilton). A gradient of water/acetonitrile/0.1% formic acid was employed for chromatography. The samples were injected into the column and run for 180 minutes at a flow rate of 0.60 μl/min. The peptide separation began with 1% acetonitrile, increasing to 3% in the first 4 minutes, followed by a linear gradient from 3% to 23% acetonitrile over 86 minutes, then another increase from 24% to 80% acetonitrile over 35 minutes, and finally a 35-minute wash at 80% acetonitrile, and then decreasing to 1% acetonitrile for 10 min and kept 1% acetonitrile for another 10 min. The eluted peptides were ionized using positive nanoelectrospray ionization (NSI) and directly introduced into the mass spectrometer with an ion source temperature set at 250°C and an ion spray voltage of 2.1 kV. Full-scan MS spectra (m/z 350–2000) were captured in Orbitrap Exploris at a resolution of 120,000 (m/z 400). The automatic gain control was set to 1e^6^ for full FTMS scans and 5e4 for MS/MS scans. Ions with intensities above 1500 counts underwent fragmentation via nanoelectrospray ionization (NSI) in the linear ion trap. The top 15 most intense ions with charge states of ≥2 were sequentially isolated and fragmented using normalized collision energy of 30%, activation Q of 0.250, and an activation time of 10 ms. Ions selected for MS/MS were excluded from further selection for 3 seconds. The Orbitrap Exploris mass spectrometer was operated using Thermo XCalibur software.

### 2.7 Data Analysis in MaxQuant

MaxQuant search engine (version 2.4.1.0) was used to process the raw mass spectrometric files [34]. The MS/MS spectra were searched using the Andromeda search engine against the UniProt-mouse protein database (UP000000589, December 2021, 55,086 entries) concatenated with reverse decoy database. Trypsin/P was used as the cleavage enzyme allowing up to two missing cleavages. Precursor mass and fragment mass were set with initial mass tolerances of 20 ppm for both the precursor and fragment ions. The carbamidomethylation of cysteine was set as a fixed modification. The oxidation of methionine and deamidation of asparagine/glutamine were set as variable modifications. The maximum missed cleavage sites were set to 2, and the maximum modifications/peptide and minimum peptide length were set at 6 amino acids. The false discovery rate (FDR) of peptide spectrum match (PSM) and protein identifications was set at 1%, which was calculated from the target-decoy search approach. For relative quantification, the MaxQuant’s label-free quantification function, LFQ and iBAQ were enabled [35].

### 2.8 Statistical analysis

Data were collected from three biological replicates for each organ. The protein groups.txt output from MaxQuant was exported into Perseus v1.6.15.0 for downstream analysis [36]. The protein groups identified from the reverse decoy database, or marked as potential contaminants, or only identified by a post-translational modification site were removed. The remaining intensity, label-free quantitation (LFQ) intensity and iBAQ intensity values were log_2_ normalized. For purified library replicates, an LFQ analysis was performed to assess the reproducibility with a Pearson correlation coefficient. To find the differential abundance of proteins, a student’s t-test was conducted with an artificial within-groups variance, s0, set at 0.1. The test was applied on data filtered for only those proteins with a valid LFQ intensity in at least all replicates in each of the mouse organ sample preparations. Before applying the t-test, the remaining undefined intensity values were imputed from a normal distribution with a downshift of 1.8 standard deviations from the total sample mean and a width of 0.3 times the sample standard deviation. In this study, proteins are considered differentially expressed between the normal and disease conditions or different types of tissues if the peptide intensity fold change (FC) across samples is ≥ 2 or ≤ -2 (the absolute value of log_2_FC ≥ 1), p < 0.05 (-log_10_ (p-value) > 1.3).

### 2.9 Protein annotation

The protein list obtained from MaxQuant was subjected to a gene ontology GO-term analysis using the UniProtKB database to identify proteins with the GO-term “membrane”, which were then annotated as membrane proteins (MPs). Within the MPs group, proteins containing at least one α-helical transmembrane segment were labeled as IMPs (for integral membrane proteins). The Phobius web server was utilized (http://phobius.sbc.su.se/) to predict the number of transmembrane segments (TMS). The subcellular localization of the IMPs was further classified using the GO-term “Subcellular location [CC].” These were divided into pIMPs (for plasma integral membrane proteins) if they contain the GO-term ‘plasma membrane’. Alternatively, they were labeled as oIMPs (for organelle or other integral membrane proteins) when containing keywords such as ER, Golgi membrane, vesicle membrane (including endosome, exosome, peroxisome, lysosome, and vesicle), mitochondrial and nucleus membranes).

### 2.10 Functional enrichment analysis

The datasets were analyzed using the Database for Annotation, Visualization and Integrated Discovery (DAVID; https://david.ncifcrf.gov/) to determine enriched biological processes [37]. The Gene Ontology Biological Processes (‘GO_BP_Direct/GO_BP_FAT’) and KEGG Pathways (‘KEGG_PATHWAYS’) were used to classify the enriched functions. An Ease score was applied, which is a modified Fisher exact p-value that is more conservative. Terms with scores greater than 0.05 were considered significant. ClueGo App on Cytoscape was used to visualize the interconnection between gene ontology biological process terms for our significantly dysregulated proteins [38]. A right-sided hypergeometric test (Benjamini–Hochberg FDR <0.02) and kappa scores (κ≥0.4) determined term-term correlation.

## 3. RESULTS AND DISCUSSION

### 3.1 Identification of the mouse organ membrane proteome using peptidisc and SP4 methods

We recently reported the characterization of the mouse organ membrane proteome using the peptidisc method [39]. Here, we evaluated the method’s efficiency compared to the recently introduced SP4 method [40]. The SP4 protocol entails the capture of acetonitrile-induced protein aggregates through a centrifugation process using glass beads. This technique has been acknowledged for its ability to isolate low-solubility aggregates that often include insoluble transmembrane proteins [40]. On the other hand, the peptidisc captures transmembrane proteins in a soluble state using a His_6_-tagged scaffold that enables their selective recovery through Ni-NTA chromatography [41,42]. Despite the distinct approaches, the methods share a common initial step of ultracentrifugation to enrich the membrane fraction and deplete high-abundance soluble proteins that often obscure membrane protein detection [43]. To benchmark our analysis, we employed two mouse organs, liver and lung, each in three biological replicates. As summarized in **Table 1 (**from datafile **Table S1)**, the peptidisc method identified a total of 1,508 and 2,200 proteins in the liver and lung samples, respectively, with approximately 43% of these proteins classified as integral membrane proteins (hereafter termed **IMPs**). The SP4 method detected 2,200 and 3,050 proteins in the liver and lung samples (**Table 1** and **Table S2**), respectively, with approximately 27% annotated as IMPs. Thus, the SP4 method detected a higher number of proteins overall but retrieved slightly fewer IMPs compared to peptidisc.

**TABLE 1.**
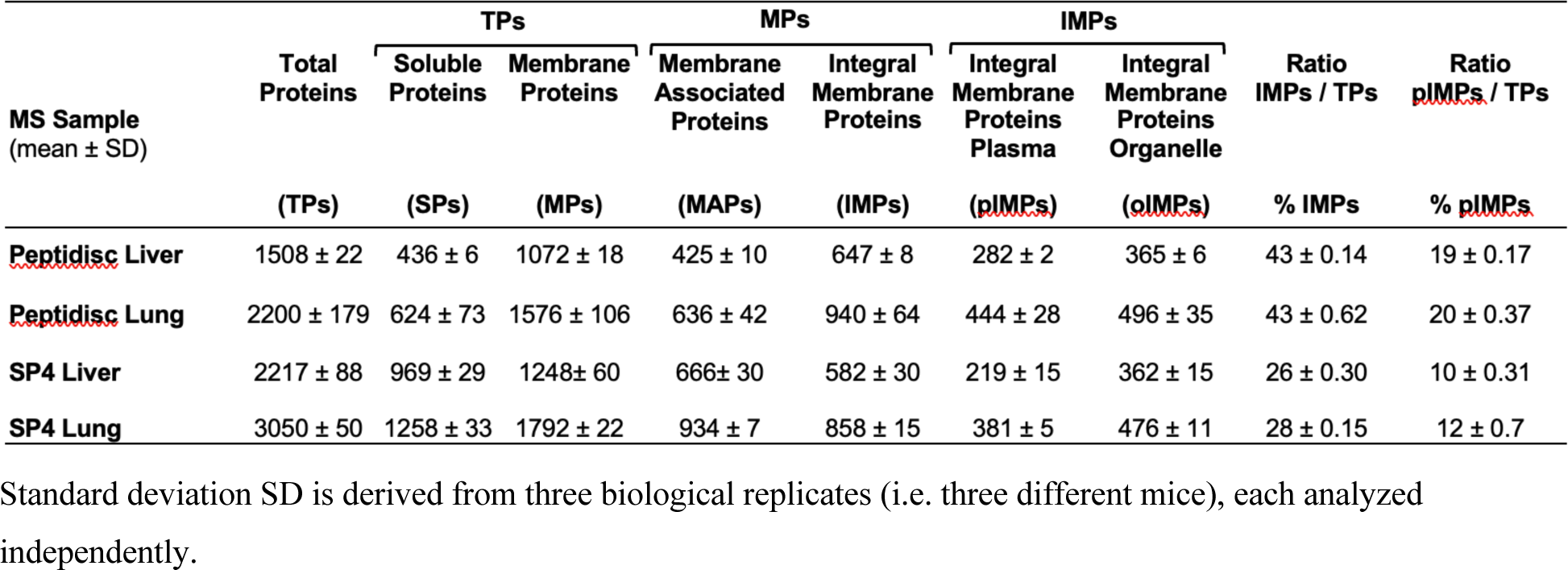
Number and type of proteins identified in the peptidisc and SP4 methods.

### 3.2 Performance of the peptidisc and SP4 methods in identifying differentially expressed integral membrane proteins

As an initial assessment of the method performances, we determined the IMPs’ relative abundance over total proteins (TPs) by plotting their intensity-based absolute quantification values (iBAQ; **Figure 1A** and **Table S3**). Remarkably, the peptidisc method showed a higher representation of IMPs in the first and second quartiles compared to SP4, probably due to the more effective removal of soluble protein contaminants. We then determined the methods’s efficacy in capturing liver-specific IMPs. As we reported recently, a protein was considered to be liver-specific (*i.e.* organ enriched) when its relative abundance is heightened at least 2.5-fold (Log_2_) compared to another organ, here the lung [39]. Through this assessment, the peptidisc identified 146 liver-specific proteins of which 80 were classified as IMPs (**Figure 1B** and **Table S1**). The SP4 method identified 288 liver-specific proteins of which 40 were classified as IMPs (**Figure 1B** and **Table S2**).

**FIGURE 1.**
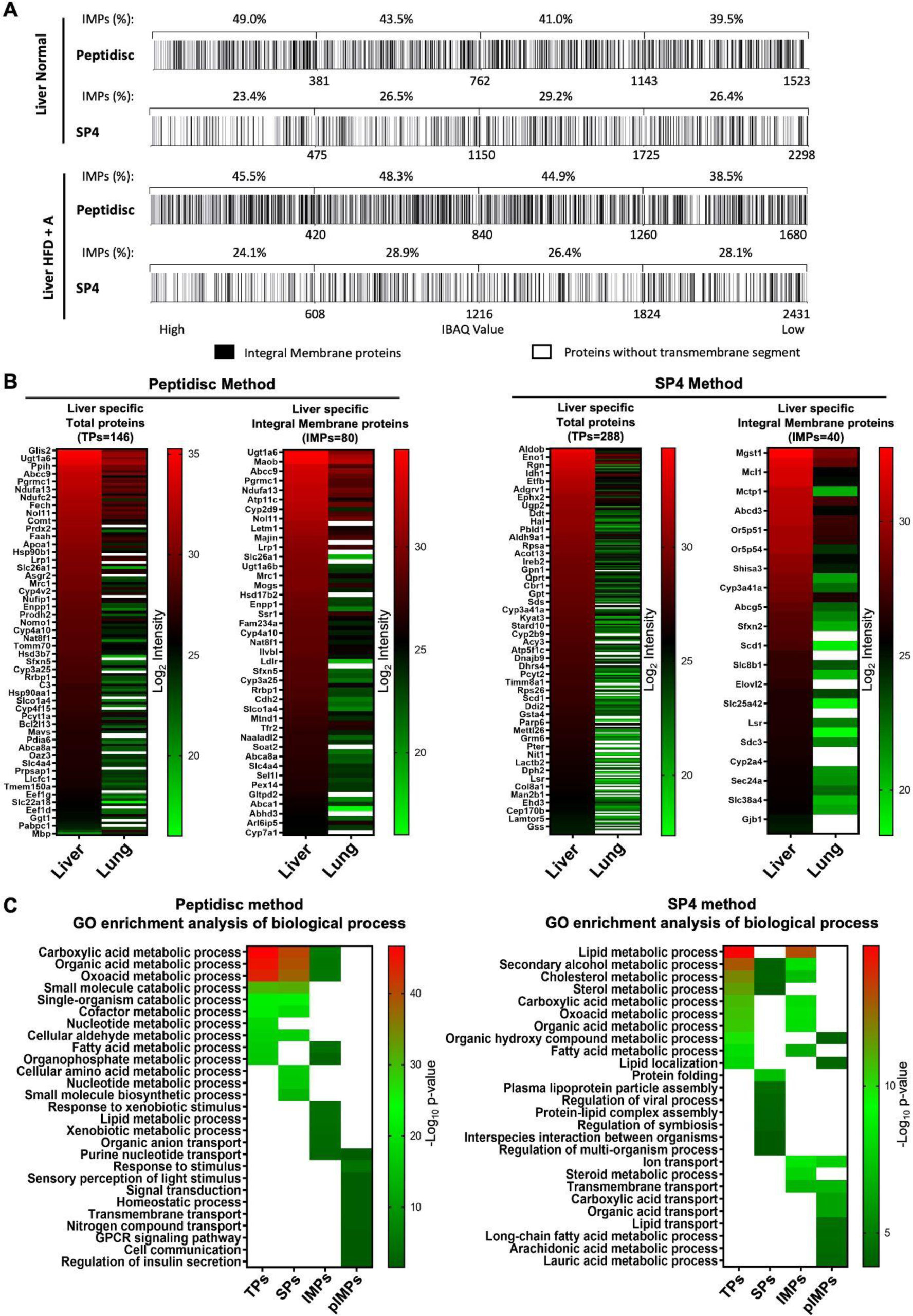
Comparing peptidisc and SP4 methods. (**A**) Abundance of integral membrane proteins (IMPs, black bars) relative to soluble proteins (white bars). The total proteins (TPs) identified in each organ are ranked according to their iBAQ value and plotted on the graph divided into quartiles. The relative abundance of IMPs in each quartile is provided as a percentage of TPs. (**B**) Abundance heatmap of liver-specific proteins. Liver-specificity was arbitrarily defined when the protein abundance in the liver sample is ≥ 2.5-fold (log_2_) compared to the lung sample, provided its p-value is < 0.05. The peptide intensity is given in a color scale from green (low) to red (high). Due to space limitation, the TPs map displays every third gene name only. The white field color indicates that the protein is not detected. **(C)** Gene ontology enrichment analysis of biological processes (‘GO_BP_Direct’/ ‘GO_BP_FAT’) of liver-specific proteins. The top 10 biological processes associated with the liver-specific total proteins (TPs), soluble proteins (SPs), integral membrane proteins (IMPs) and plasma membrane integral membrane proteins (pIMPs) are presented. The scale is based on log_10_ (p-value), with a higher value (red) indicating stronger evidence for the correlation between the dataset and the function. The white field color indicates that the corresponding biological process is not assigned to the sample. Log_10_ (p-value) > 1.3 is considered statistically significant. Produced with the DAVID bioinformatics resources.

We then utilized the DAVID bioinformatics resources to investigate the biological processes tied to the liver-specific dataset [37]. As shown in **Figure 1C**, the top 10 biological processes linked to the liver-specific dataset aligned remarkably well with the anticipated liver functions; both methods presented a strong correlation to processes such as lipid transport, cholesterol, organic acid and alcohol metabolism [44]. When narrowing the analysis to the soluble protein (SPs) dataset, the SP4 method still captured the liver-specific functions, but the peptidic showed less correlation (**Figure 1C**). Interestingly, when inspecting the plasma membrane-located IMPs (hereafter termed pIMPs) dataset, the SP4 method could no longer demonstrate this correlation. For example, liver-specific pIMPs were linked with ‘sensory perception of light stimuli,’ whereas the peptidisc continued to capture with high confidence lipid-related functions (**Figure 1C**). Consequently, when considering the membrane proteome, the SP4 method was less effective than the peptidisc method in identifying proteins that reflect the liver organ specificity. Further research is required to explain this difference, but presumably, the larger “spectral space” due to the larger soluble proteins pool in the SP4 method (**Figure 1A**) is likely to decrease the detection accuracy for the less abundant IMPs.

### 3.3 Identification of dysregulated IMPs in the diseased mouse liver

Since the peptidisc performance was found adequate to identify the liver proteome functions at the membrane protein level, we extended the method to explore the disparity between normal and disease states, here induced through a combination of high-fat diet and alcohol consumption (termed HFD+A hereafter). The liver, and as a control the lungs, were obtained post-treatment and processed to capture the membrane proteome in peptidiscs. The number of total proteins (TPs) identified in the treated mice samples was comparable to the untreated mice, with approximately 43-46% classified as IMPs (**Table 2**). The peptide intensity value for the commonly identified proteins was then plotted to help visualize the level of dysregulation (**Figure 2A**). The plots revealed a notable difference for the liver proteome, whereas the lungs were minimally affected by the treatment (correlation coefficients of 0.83 and 0.94, respectively). Noteworthy, only five IMPs exhibited a two-fold expression change (Log_2_) in the lungs, contrasting to 117 IMPs in the liver. The obvious level of dysregulation of the liver sample is consistent with the HFD+A treatment that primarily targets the liver and much less the lungs.

**FIGURE 2.**
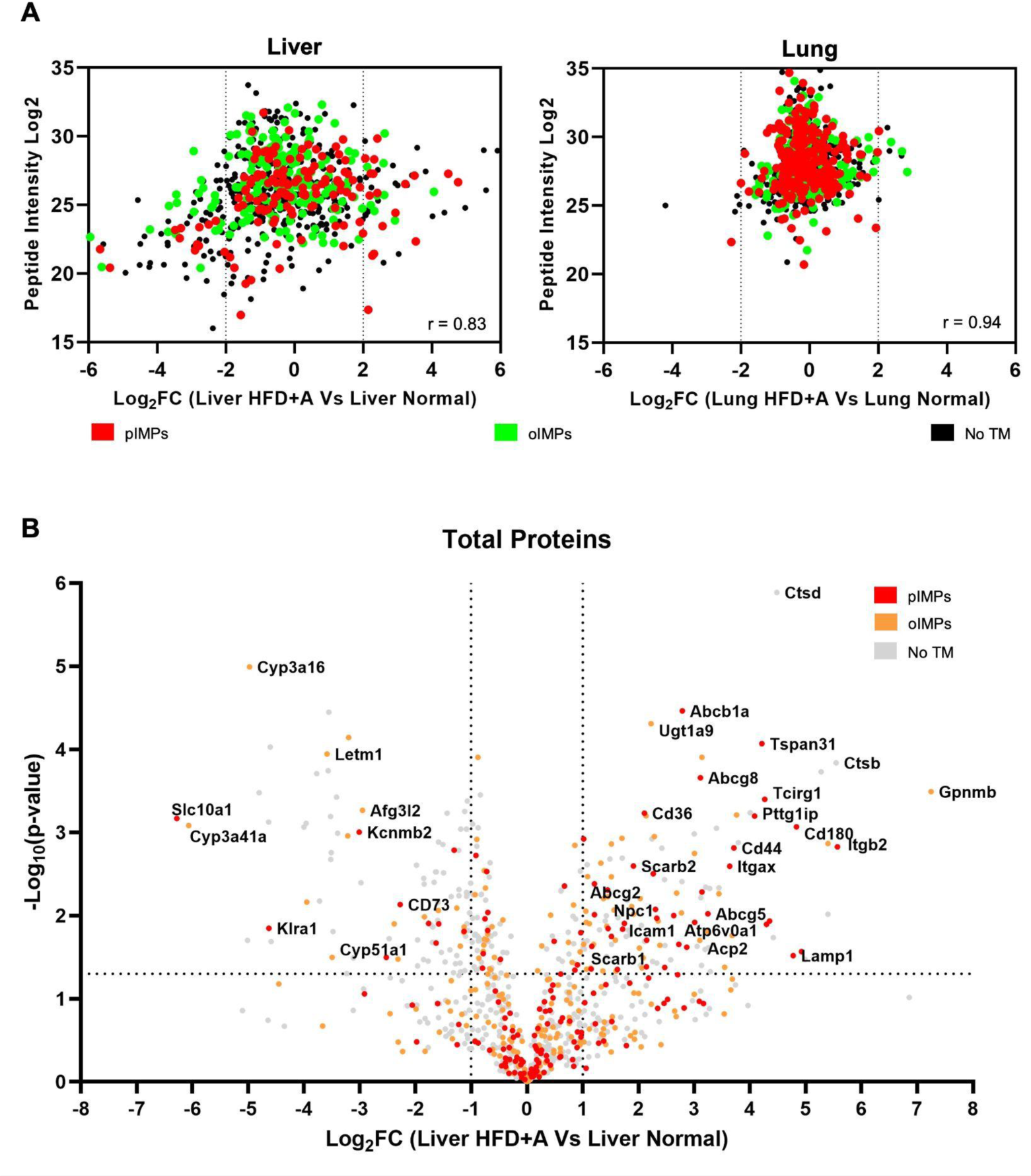
Dysregulated proteins after the HFD+A treatment. (**A**) Scatter plot depicting peptide intensity variation between treated and untreated mice, highlighting differentially expressed proteins in the liver and lungs on one replicate. Proteins without transmembrane segments (no TM) are colored in black. The dotted lines indicate a 2-fold change cut-off (Log_2_FC ≥ 1). The Pearson correlation coefficient (r) was obtained by using MaxQuant and Perseus software (**B**) Volcano plot representation of dysregulated proteins in biological replicates (n=3). The significance (non-adjusted p-value) and the fold-change were converted to −Log_10_ (p-value) and Log_2_ FC, respectively. The vertical and horizontal dotted lines show a cut-off value of Log_2_ FC ≥ ±1 and of Log_10_ (p-value) ≤ 1.3, respectively. The volcano plot was generated using Perseus software and Graph Pad Prism. Proteins without transmembrane segments (no TM) are colored gray.

**TABLE 2.** Number and type of proteins identified in the liver and lungs of treated and untreated mice using the peptidisc method.

| MS Sample<br>(mean ± SD) | TPs | | | MPs | | IMPs | | Ratio<br>IMPs / TPs | Ratio<br>pIMPs / TPs |
| --- | --- | --- | --- | --- | --- | --- | --- | --- | --- |
|  | Total<br>Proteins | Soluble<br>Proteins | Membrane<br>Proteins | Membrane<br>Associated<br>Proteins | Integral<br>Membrane<br>Proteins | Integral<br>Membrane<br>Proteins<br>Plasma | Integral<br>Membrane<br>Proteins<br>Organelle |  |  |
|  | (TPs) | (SPs) | (MPs) | (MAPs) | (IMPs) | (pIMPs) | (oIMPs) |  |  |
| Liver Normal | 1508 ± 22 | 436 ± 6 | 1072 ± 18 | 425 ± 10 | 647 ± 8 | 282 ± 2 | 365 ± 6 | 43 ± 0.14 | 19 ± 0.17 |
| Liver HFD+A | 1563 ± 89 | 415 ± 30 | 1148 ± 59 | 433 ± 21 | 715 ± 38 | 317 ± 20 | 397 ± 18 | 46 ± 0.16 | 20 ± 0.16 |
| Lung Normal | 1715 | 392 | 1323 | 492 | 831 | 416 | 415 | 48 | 24 |
| Lung HFD+A | 1617 | 371 | 1246 | 469 | 777 | 378 | 399 | 48 | 23 |

The standard deviation SD is derived from three biological replicates for the liver sample. The determination of the SD for the lung sample was not deemed necessary for this analysis.

We then ranked the liver proteins identified in peptidiscs based on their fold change to obtain a list of proteins most evidently dysregulated (**Figure 2B** and **Table S4**). This list comprises proteins with a fold change greater than two (Log_2_ FC ≥ 1) and a p-value lesser than 0.05 (-Log_10_ (p-value) >1.3). Using these criteria, 128 proteins (∼8.5% of the dataset) were upregulated after the HFD+A treatment and 93 proteins (∼6.2% of the dataset) were downregulated. Among the 221 dysregulated proteins, 106 were classified as IMPs, including 48 pIMPs and 58 oIMPs (IMPs located in organelle membranes). We also applied the SP4 method to the same liver membrane preparations, but a lower number of proteins was found dysregulated using this method (76 proteins including 25 IMPs; **Figure S1** and **Table S5**). This later observation further validates the utility of the peptidisc in identifying MP dysregulation in the context of organ disease.

### 3.4 Dataset analysis using the DAVID, ClueGO and KEGG bioinformatics resources

Before commenting on the dysregulated IMPs, we determined the main biological processes affected by the HFD+A treatment. This was accomplished using Gene Ontology (GO) enrichment analysis with the DAVID bioinformatics resources (**Figure 3A**) [37]. The results revealed that many enriched functions are related to lipid processing, including lipid metabolism, transport and cholesterol homeostasis. We then employed ClueGO to visualize the biological network formed by the upregulated IMPs, with term-term interaction based on shared genes between the terms (**Figure 3C**) [38]. The interconnections showed that a majority of upregulated proteins are related to lipid-related biological processes. Finally, we employed the Kyoto Encyclopedia of Genes and Genomes (KEGG) to name the pathways principally affected (**Figure 3B**). The results pointed to cell adhesion, cholesterol metabolism, bile secretion and lysosomal processes as the main pathways augmented by the HFD+A treatment. The upregulation of these pathways was somewhat expected given the HF diet and alcohol promote fat-liver accumulation [45,46].

**FIGURE 3.**
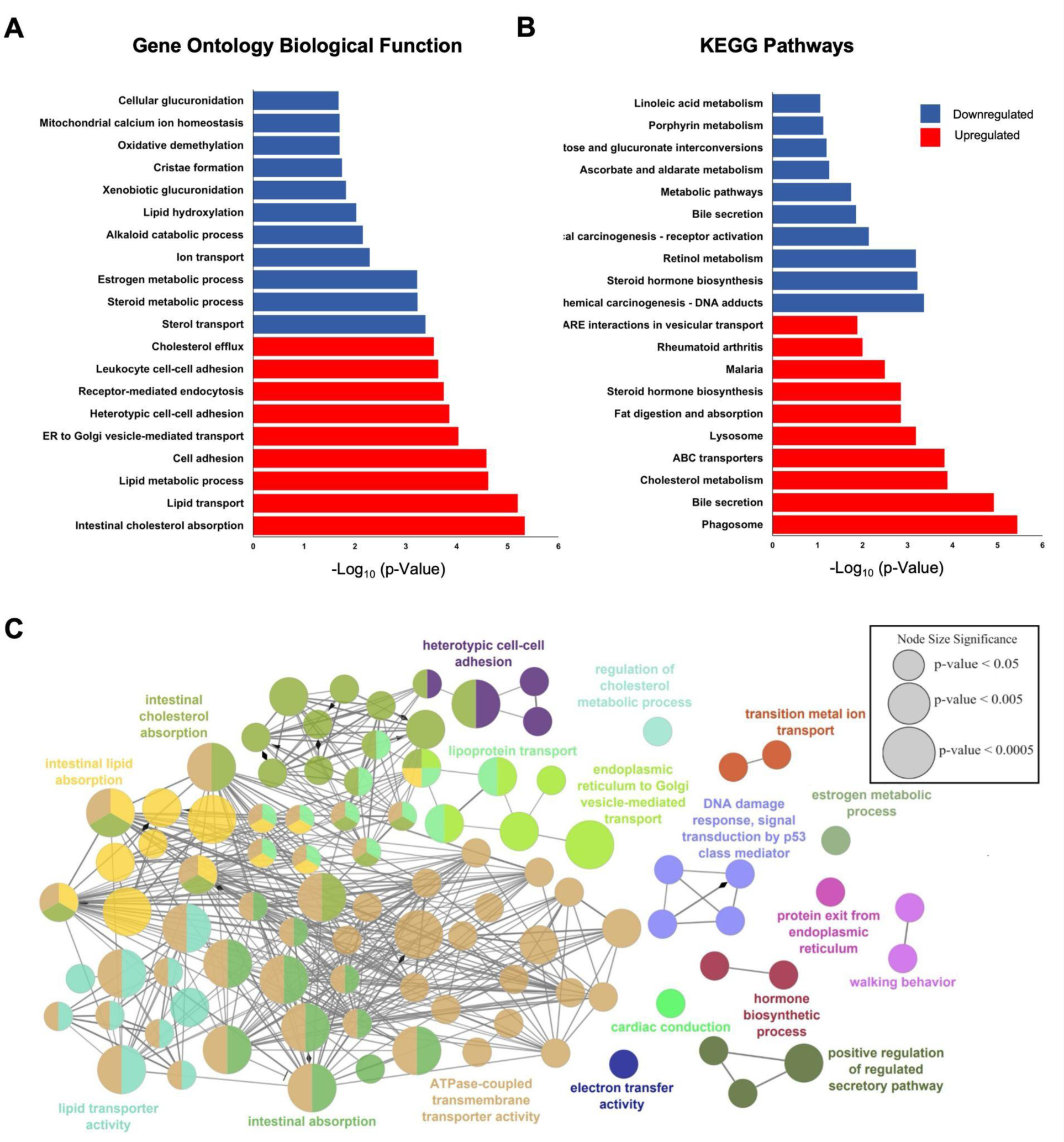
Bioinformatic analysis of dysregulated proteins. (**A**) Gene Ontology (GO) enrichment analysis of biological processes for downregulated (blue) and upregulated (red) IMPs after the HFD+A treatment. The top 10 biological processes are presented and the scale is based on Log_10_ (p-value), with a value greater than 1.3 considered statistically significant. Produced with the DAVID Bioinformatics Resources. **(B)** Top 10 enriched Kyoto Encyclopedia of Genes and Genomes (KEGG) pathways of upregulated and downregulated IMPs after the HFD+A treatment. **(C)** ClueGO network analysis of upregulated proteins. GO biological processes terms are linked based on their kappa score (≥0.4). Functionally related terms are grouped and color coded based on overlapping proteins. Only the most significant terms in each group are shown. Node size represents the term enrichment significance.

### 3.5 Dysregulation of IMPs involved in cholesterol processing

According to the global analysis above, six membrane proteins associated with cholesterol and triglycerides metabolism showed upregulation (i.e. SCARB1, CD36, NPC1, ABCG5, ABCG8, and LRP1). SCARB1 aids in liver uptake of cholesterol esters from circulating HDL, whereas ABCG5/G8 promotes the excretion of cholesterol to the bile ducts [47,48]. We also note that a set of lysosomal IMPs such as NPC1, SCARB2, ATP6V0A1, TCIRG1, ACP2 and LAMP1 were upregulated, whereas the ER membrane Cyp51a1, which contributes to cholesterol de novo synthesis [49], was downregulated (**Figure 2B**). The lysosome is the main site for processing cholesterol esters, and NPC1 and SCARB2 directly contribute to cholesterol transport from the lysosomal interior to the cell cytosol for excretion [50]. Thus, collectively, the dysregulation of the IMPs would augment hepatic cholesterol uptake, increase its degradation and excretion, and decrease de novo synthesis. These joint regulations would result in enhanced cholesterol elimination, presumably as a protective mechanism against the detrimental effects of a high-fat diet and alcohol exposure.

### 3.6 Dysregulation of IMPs involved in bile salts excretion and reabsorption

Bile-acid synthesis is a main pathway for cholesterol catabolism, and several IMPs related to bile salts processing were dysregulated in our dataset (i.e. SLC10A1, ABCB1A, ABCG2, UGTlA5, UGT1A9, UGT2B1 and UGT2B35) [51]. Particularly noteworthy is the 6.3-fold (Log_2_) reduction in the expression of SLC10A1 (**Figure 2B)**. This solute carrier promotes the reabsorption of conjugated bile acids from the hepatic portal vein as part of the enterohepatic circulation [52]. The downregulation of SLC10A1 may thus trigger a feedforward upregulation of hepatic bile-acid synthesis from hepatic cholesterol, leading to a decrease in circulating sterol levels. This is in line with the observation that patients with ALD and NAFLD often exhibit bile acidemia and elevated bile-acid urinary excretion [53-55]. Furthermore, SLC10A1 knockout mice are better protected against NAFLD, showing the benefits of reducing bile acid reabsorption [56]. We also note the upregulation of ABCB1A. This multidrug transporter exports substances like conjugated bile acids into the bile ducts and its upregulation upon exposure to xenobiotics such as ethanol has been documented [57-59].

### 3.7 Dysregulation of xenobiotic metabolic processes

The UDP-glucuronosyltransferase (UGT) family plays a major role in the detoxification of xenobiotics and the conversion of bilirubin to a water-soluble form eliminated through bile or urine. Studies have shown a strong relationship between low Ugt1a activity, bilirubin metabolism disorders, and increased liver damage [60]. Here, we speculate that the observed downregulation of Ugt1a5 may cause the upregulation of the closely related Ugt1a9 in our dataset, perhaps as a compensatory response. Alternatively, since Ugt1a9 is one of the primary enzymes used for ethanol glucuronidation [61], its upregulation could be directly related to alcohol treatment. Our dataset also shows dysregulation of CYP3A16 and CYP3A41a enzymes. The cytochrome P450 proteins catalyze many redox reactions, and although the exact relationship between CYP3A16, CYP3A41a, and liver disease is less documented, their reduced expression has been observed in NAFLD [62, 63]. Of particular interest, Cyp3a gene knockout in mice leads to increased bile acid synthesis from cholesterol, raising the intriguing question of whether the downregulation of CYP3A16 and CYP3A41a might also influence the bile secretion pathway in the treated mouse model [64].

### 3.8 Cell adhesion and mitochondrial dysfunction

Given the central role of cell adhesion molecules in the inflammatory response, it is no surprise that ITGAX (Cd11c) and ITGB2 (Cd18) are upregulated in our dataset. These proteins combine to form the CR4 receptor, which in mice is primarily expressed on dendritic cells to perform cell adhesion, phagocytosis, and migration [65,66]. This receptor recognizes different cell surface ligands including ICAM-1 (CD54) which is also upregulated [67]. Thus, the higher expression of these proteins together may support an increased mobility of dendritic cells to damaged liver tissue, thereby supporting the tissue inflammatory response [68]. Mitochondria damage, in addition to cholesterol accumulation, may be at the origin of this inflammation. Mitochondrial defects, especially in cristae formation, disrupt the membrane proton gradient as previously noted in ALD and NAFLD [22-24,69]. Here we identified two down-regulated proteins, coded by AFG3L2 and LETM1, as potential contributors to mitochondrial abnormalities. The LETM1 gene encodes a proton/calcium antiporter crucial for maintaining the tubular shape of mitochondria, and AFG3L2 is a protease with a critical role in proteostasis and cristae morphogenesis [70,71]. Further investigation into these protein functions could offer insights into mitochondrial dysfunction in ALD and NAFLD.

### 3.9 GPNMB and CD44

Finally, given the massive upregulation of GPNMB (7.2-Log_2_ equivalent to ∼160 fold), we delved into the role of this single-pass transmembrane glycoprotein, in conjunction with the CD44 receptor also upregulated in the dataset. In hepatocytes, a fraction of Gpnmb is cleaved with ADAM10 to become secreted [72]. In its soluble form, Gpnmb binds to adipocytes via the CD44 receptor to promote lipogenesis [73]. Considering that the HF diet and alcohol shift the hepatocyte’s metabolic pathways from oxidative to reductive synthesis causing fatty acid synthesis, the observed upregulation is logical [74,75]. We note, however, that in macrophages Gpnmb binding to CD44 instead inhibits the NF-κB signaling pathway to decrease the inflammatory response [76]. The macrophage-produced Gpnmb also promotes the proliferation of mesenchymal stem cells, which are typically recruited for repairing tissue damage [77]. This is in line with the role of CD44 on hepatic progenitor cells which promotes liver regeneration through increased uptake of extracellular cystine [78]. Further research is thus warranted to elucidate the precise mechanisms involved and to determine which cells are responsible for the increased expression of GPNMB and CD44. Interestingly, CD44 integrates into tetraspanin-enriched microdomains in the plasma membrane [79]. This preferential interaction may explain why TSPAN31 is notably upregulated in the treated sample. Similarly, CD36 in many cells is localized in specialized cholesterol-rich membrane microdomains [80]. We thus speculate that these tetraspanins help to stabilize the upregulation of these proteins in the plasma membrane.

## 4.0 CONCLUDING REMARKS

This study explores the membrane proteome of a diseased liver to obtain a global understanding of its dysregulation at the level of its constitutive transporters, receptors, channels and membrane enzymes among others. Our diseased mouse model was expected to mimic a realistic Western scenario involving alcohol consumption and HF diet [81]. To start this analysis, we compared the peptidisc and SP4 methods to find that peptidisc adequately profiles the liver-specific integral membrane proteins (IMPs). The SP4 method was found effective at capturing liver-specific soluble proteins, but at the same time these proteins seemingly overshadowed IMP detection, decreasing their relative quantitation. The peptidisc method instead enriched the IMPs population by depleting soluble proteins, and therefore the method emerged as more suitable for this analysis. We acknowledge that peptidisc and SP4 are not the only methods available for identifying membrane proteins and that various enrichment strategies and detergent depletion strategies exist, warranting further work to determine the optimal approach for quantifying MPs at the organ level [82].

Overall, the peptidisc successfully identified IMPs previously related to liver dysfunction (see **Table S6** for the relationship to other studies), confirming the ability of this method to capture information relevant to the disease condition at the global membrane proteome level. Analysis of the dysregulated IMPs from the liver organ revealed their predominant association with cholesterol and bile acid metabolism. We observed upregulation of IMPs involved in cholesterol uptake and secretion of bile salts, alongside a downregulation of IMPs related to bile acid reuptake and cholesterol biosynthesis. This coordinated membrane proteome response would serve to lower the reuptake of conjugated bile acids while increasing the excretion of bile components to eventually decrease harmful bile acid and cholesterol accumulation within hepatocytes. We also observed dysregulation of proteins involved with the mitochondrial membrane structure and upregulation of membrane receptors involved in the mobility of immune cells, likely contributing to the immune response of the diseased tissue. Targeting these cell adhesion molecules may prove helpful in mitigating the inflammation associated with this disease.

While providing valuable insights into the membrane proteome variations, a few limitations should be noted. For instance, understanding the exact role of individual MPs can be challenging, particularly with multifunctional proteins like CD44, which exhibit diverse functions across cell types and isoform variants. Further proteomic analysis focusing on specific cell or tissue populations or isoforms could provide deeper insights into liver dysregulations. Our proteomic analysis was also performed on diseased organs mimicking end-stage liver failure. However, it has been documented that dysregulated protein expression is non-uniform throughout treatment [83]. This temporal variability presents a challenge in accurately assessing the immediate and long-term effects of treatment on the liver. Without a comprehensive time point analysis, distinguishing between the immediate treatment effects and the liver’s adaptive changes becomes difficult. Furthermore, while this proteomic analysis on mice enhances our understanding of human disorders, it is important to note that this model does not fully replicate the physiological or pathological changes seen in the human liver [84]. Finally, different treatment types, although mimicking the same liver disease, can have different modes of dysregulation [85]. For example, genetically induced obesity and HF diet-induced obesity showed different dysregulations, leading to different causes for liver disease symptoms [86].

With these limitations in mind, overall, our results still provide a comprehensive view of the dysregulated biological processes and pathways in liver disease induced by ethanol consumption and a high-fat diet, offering a novel framework and preliminary results that contextualize the role of the membrane proteome in liver disease like never before.

## Abbreviations

ALD: alcoholic liver disease
NAFLD: non-alcoholic fatty liver disease
SP4: solvent precipitation SP3 (SP4) method
HF: high-fat diet
HFD+A: high-fat diet and alcohol consumption
MP: membrane proteins
IMPs: integral membrane proteins
pIMPs: plasma integral membrane proteins
oIMPs: organelle or other integral membrane proteins
TPs: total proteins
DDM: Detergents n-dodecyl-β-D-maltoside
SDC: sodium deoxycholate
ACN: acetonitrile
IAA: iodoacetamide
DTT: Dithiothreitol
NSI: nanoelectrospray ionization
FDR: false discovery rate
PSM: peptide spectrum match
LFQ: label-free quantitation
iBAQ: intensity-based absolute quantification
TMS: transmembrane segments
DAVID: Database for Annotation, Visualization and Integrated Discovery
GO: Gene Ontology
KEGG: Kyoto Encyclopedia of Genes and Genomes

## ACKNOWLEDGMENTS

We gratefully acknowledge Dr. Fumio Takei from the Department of Pathology and Laboratory Medicine, University of British Columbia, for his valuable collaboration and for providing diseased and normal tissues from treated and untreated mice cohorts. The authors are thankful to Jana Hodasova from the Centre for Comparative Medicine, Animal Care Services at UBC for her help in tissue dissection. Work in the Babu lab was supported by the Canadian Institutes of Health Research (CIHR) Foundation grant FDN-154318 and work in the Duong lab was supported by the CIHR Project grant PG20R34019.

## DATA AVAILABILITY STATEMENT

The MS-based proteomics data have been deposited to the ProteomeXchange Consortium via the PRIDE partner repository and are available via ProteomeXchange with the identifier PXD050825.

---

**SUPPLEMENTAL Figure S1.**
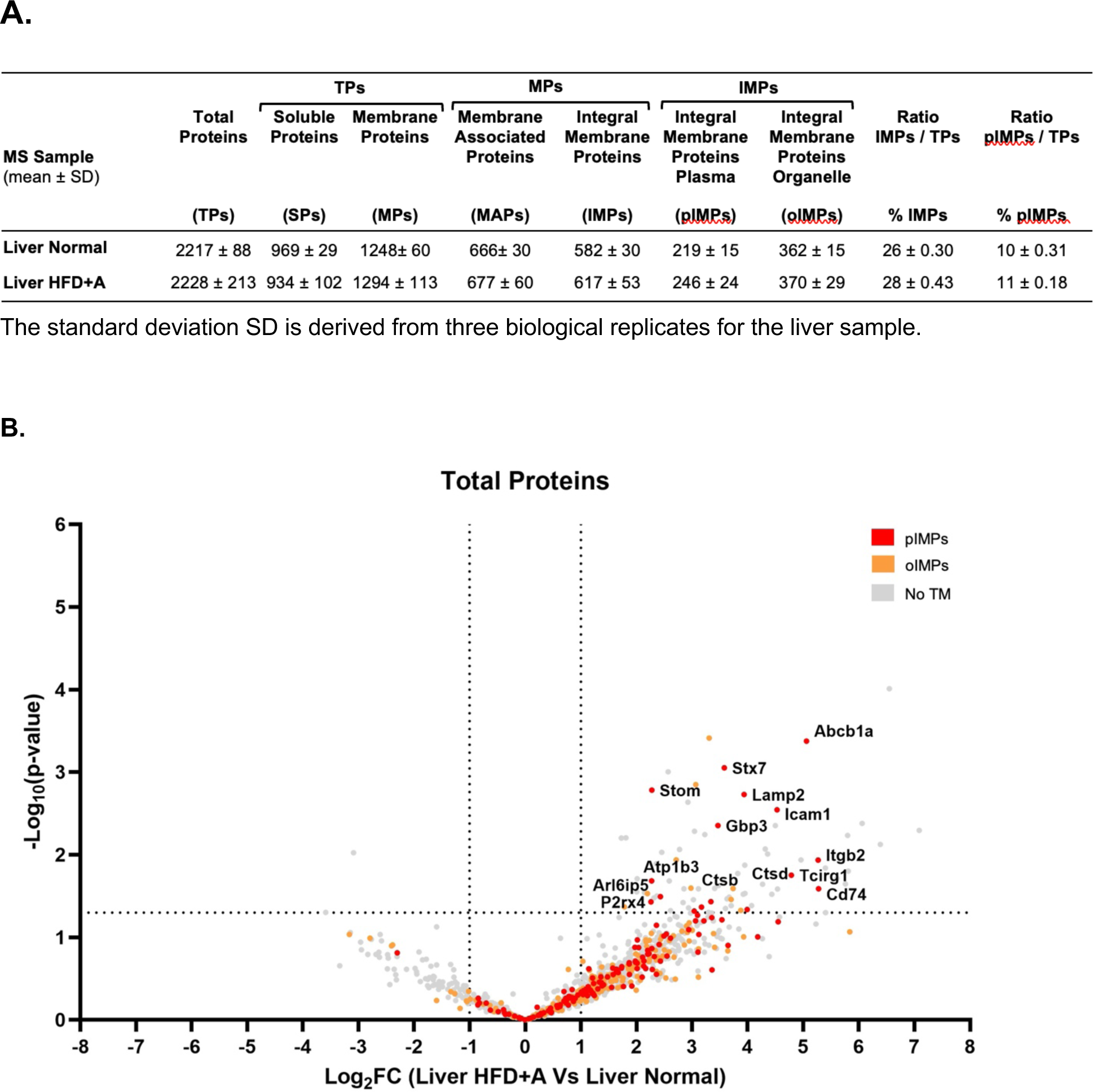
**A.** Overview of the proteins identified in the normal and treated HFD+A liver using the SP4 method. **B.** Volcano plot representation of dysregulated proteins identified using the SP4 method. The significance (non-adjusted p-value) and the fold-change were converted to −Log_10_ (p-value) and Log_2_ FC, respectively. The vertical and horizontal dotted lines show a cut-off value of Log_2_ FC ≥ ±1, and of Log_10_ (p-value) ≤ 1.3, respectively. The volcano plot was generated using Perseus software and Graph Pad Prism. Primary dataset in Table S5.

**SUPPLEMENTAL Figure S2.**
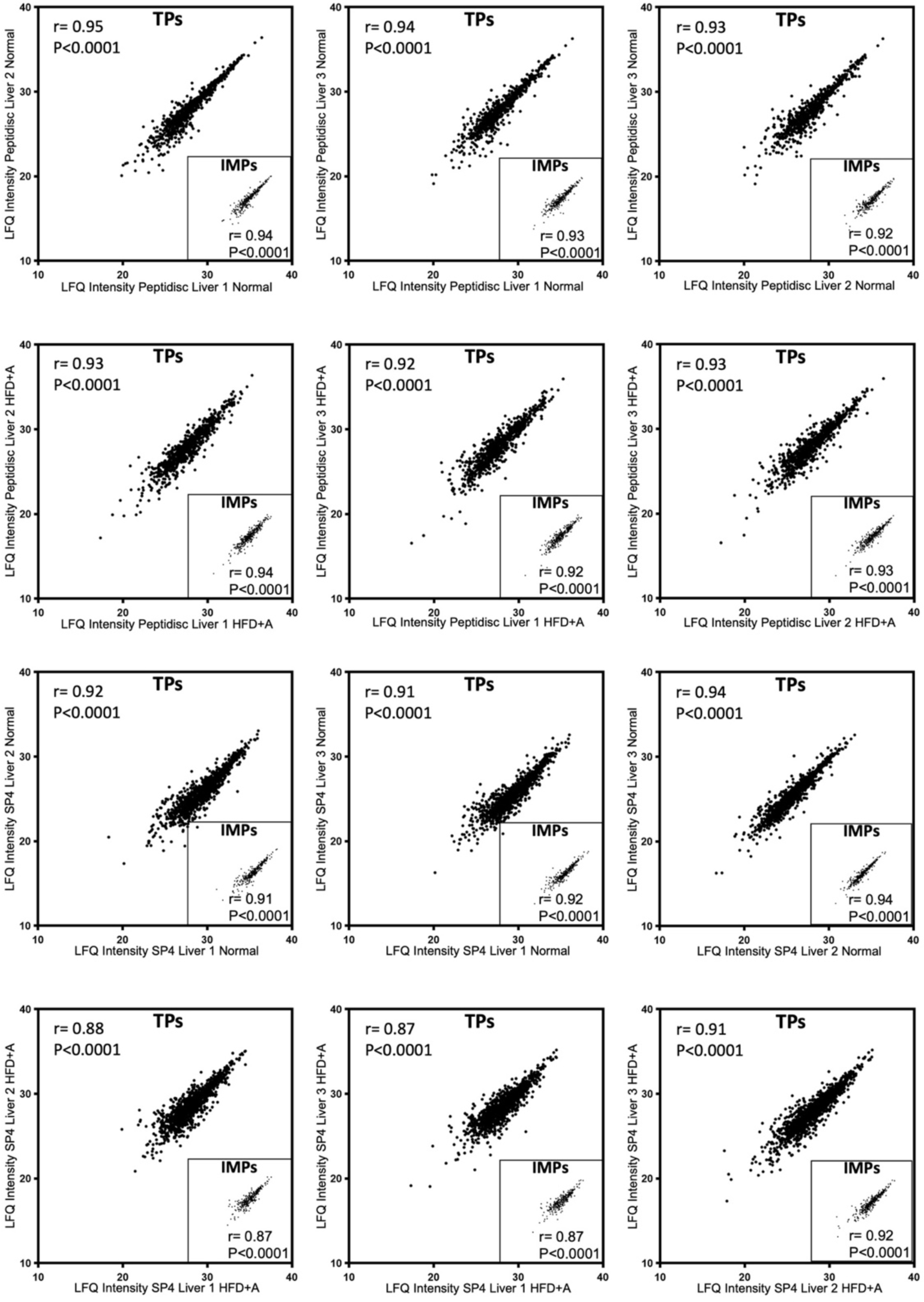
LFQ intensity variance across biological triplicates. The label-free quantitation analysis was performed using the purified libraries. The replicate comparison is presented for the liver normal or liver HFD+A. The Pearson correlation coefficient was obtained by using MaxQuant and Perseus software. The p and r values obtained for total proteins (TPs) and total integral membrane proteins (IMPs) are indicated.

**SUPPLEMENTAL Table S6.**
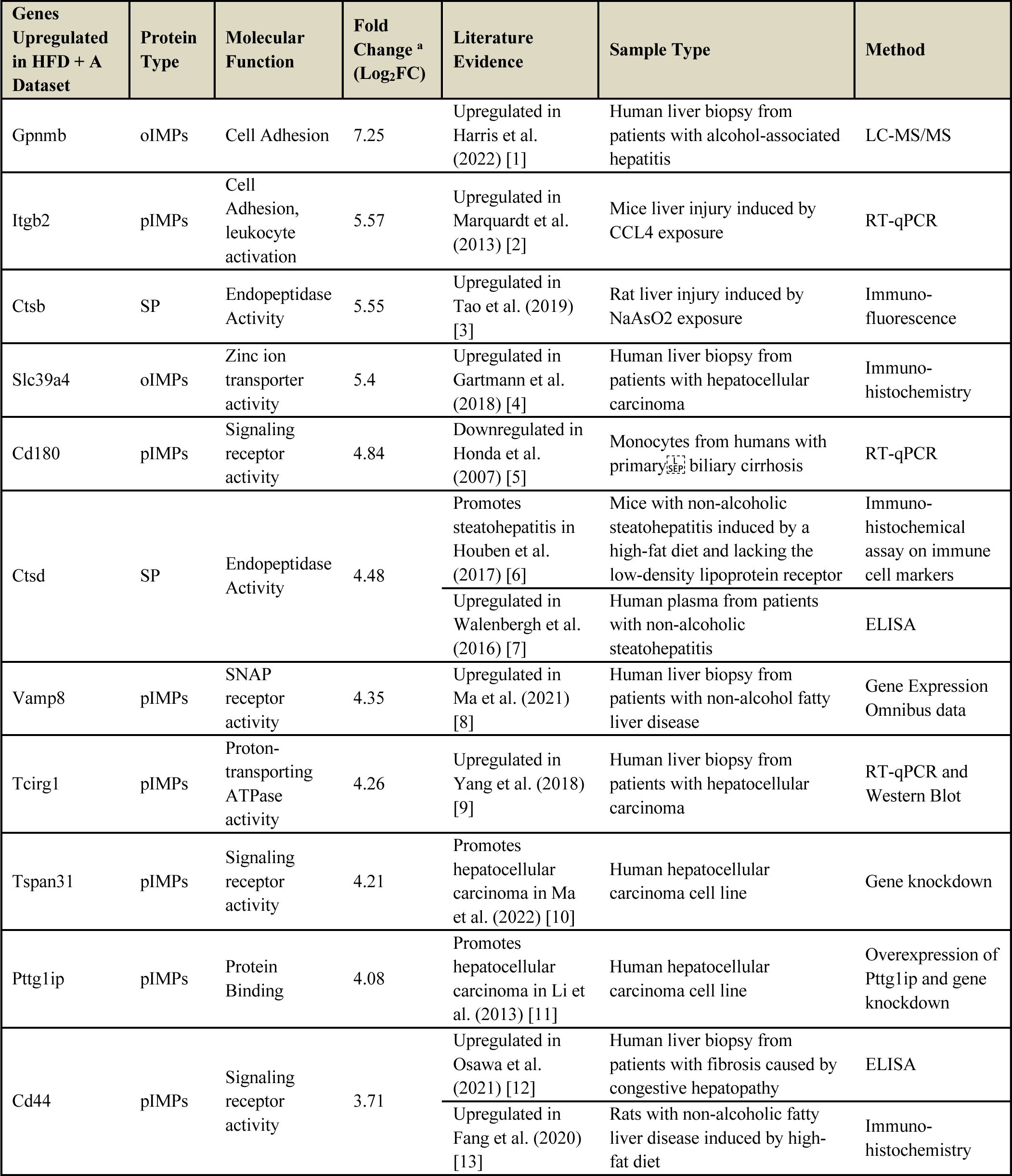
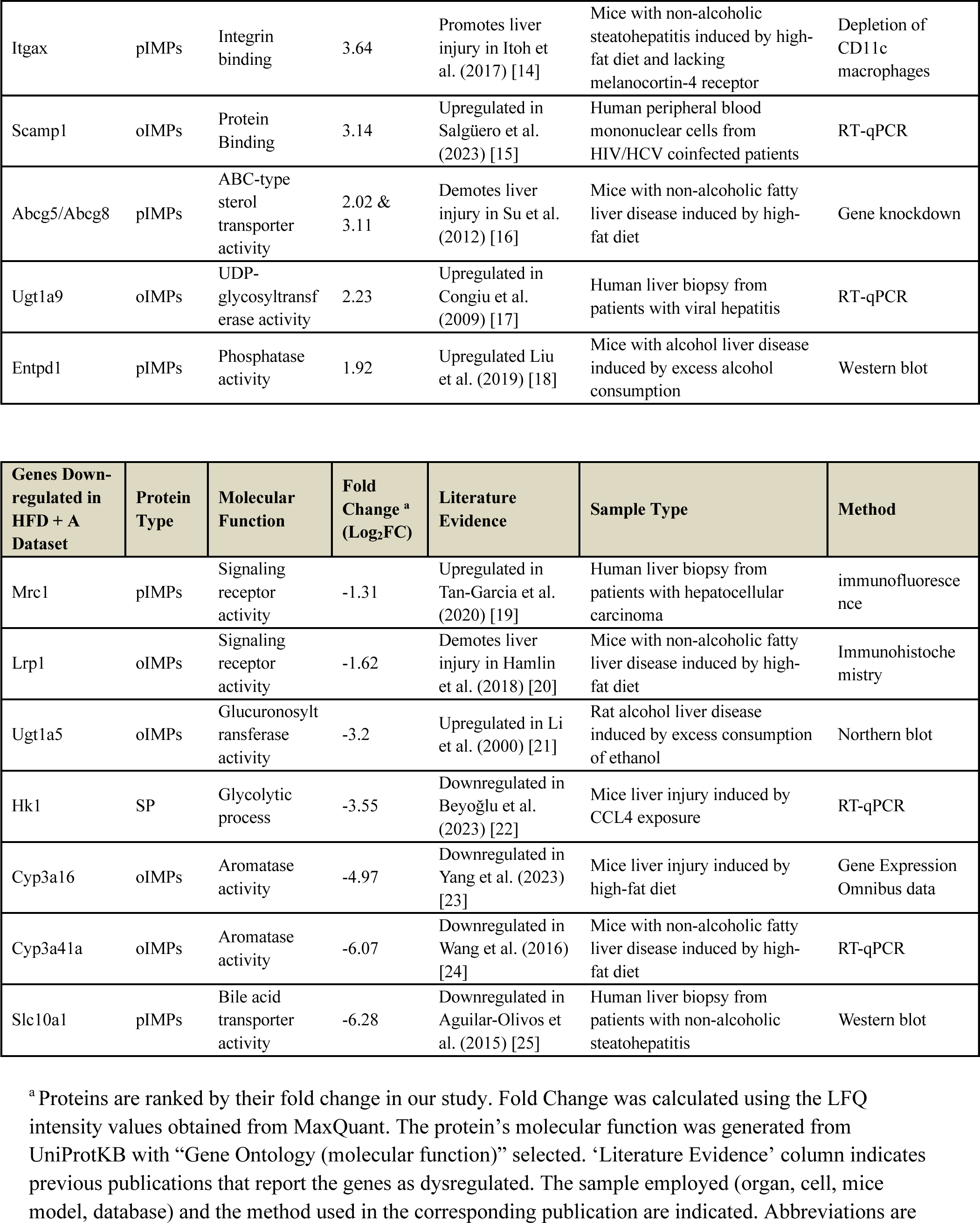
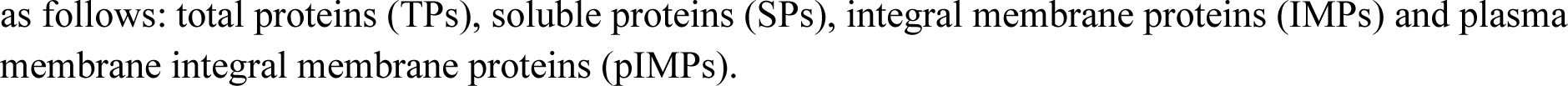
Previously established dysregulated genes in liver disease.

